# SpatialLeiden - Spatially-aware Leiden clustering

**DOI:** 10.1101/2024.08.23.609349

**Authors:** Niklas Müller-Bötticher, Shashwat Sahay, Roland Eils, Naveed Ishaque

## Abstract

Clustering can identify the natural structure that is inherent to measured data. For single-cell omics, clustering finds cells with similar molecular phenotype after which cell types are annotated. Leiden clustering is the algorithm of choice in the single-cell community. However, in the field of spatial omics, Leiden has been considered a non-spatial clustering method. Here, we show that by integrating spatial embeddings Leiden clustering is rendered into a computationally highly performant, spatially aware clustering method that compares well with state-of-the art clustering methods.

## Background

Single-cell transcriptomics has revolutionised our understanding of cellular heterogeneity by enabling the measurement of gene expression at the individual cell level. However, this high-dimensional data poses significant challenges in extracting meaningful biological insights. This can be overcome by grouping cells with similar expression profiles into distinct clusters. By partitioning cells based on transcriptional similarities, clustering facilitates the characterization of cell-type diversity within a heterogeneous cell population. Furthermore, clustering provides a basis for downstream analyses, such as differential expression, trajectory inference, and cell-cell interaction. In single-cell transcriptomics, a variety of clustering algorithms have been used and Leiden clustering has emerged as a performant choice(1). Leiden clustering can be extended to consider multiomics data via the Leiden multiplex functionality(2).

Progress in spatially resolved omics methods has empowered researchers with the ability to map gene expression in a spatial manner, transcending conventional cell clustering approaches(3). With spatial omics, scientists can discern higher-order tissue structures, termed spatial domains, by integrating spatial information alongside gene expression data. The identification of spatial domains through spatial clustering has emerged as a standard practice in constructing spatial atlases. This is instrumental in visualizing tissue anatomy, delineating tissue spatial continuity, pinpointing domain-specific marker genes, and unravelling domain-dependent molecular regulatory networks. Performance of spatial domain identification improves when leveraging the spatial information compared to non-spatial methods(4).

Leiden clustering has been typically categorised as a “non-spatial” clustering method. However, Leiden multiplex can consider spatial embeddings during clustering, thus rendering prior assumptions of the non-spatial nature of the Leiden algorithm untrue. The Leiden algorithm clusters nodes in a network by optimising a quality function, in a simple case this can be the modularity, which maximises the differences between the actual number of edges in a community and the expected number of such edges under a null model. In single-cell transcriptomics, the cells (nodes) are connected to other cells based on the distance between cells in the gene expression space (edges), usually in a dimensionality reduced latent space. Leiden multiplex enables the user to define an arbitrary number of networks (layers) with the same set of nodes that describe different modalities of edges between the nodes. So, in spatially resolved omics data, the spatial neighbourhood can be encoded by defining a spatial connectivity (based on e.g. Euclidean distance) as the weight of edges between nodes (cells or spots).

## Results and Discussion

In this study we review how Leiden clustering can utilise spatial information through selection of spatially variable genes (SVGs) instead of highly variable genes (HVGs)(5), spatially-aware dimensionality reduction through MUTLISPATI-PCA(6) (msPCA), and explicitly modelling the spatial embedding in the Leiden multiplex clustering (SpatialLeiden) (**Figure 1a**). We demonstrate their application to a 10x Visium spatial transcriptomics dataset of the mouse dorsolateral prefrontal cortex (DLPFC), the most widely used benchmark dataset for spatial clustering methods(7). This dataset consists of spatial gene expression data and histology images of 3 replicate slices from 4 donor mice, together with ground truth annotation of anatomical domains in those tissue samples. We compare the performance of a non-spatially aware and SpatialLeiden clustering to two widely used spatially-aware domain detection tools, SpaGCN and BayesSpace(8,9), and evaluating performance of the tools **(Figure 1 b, c)**.

**Figure 1:**
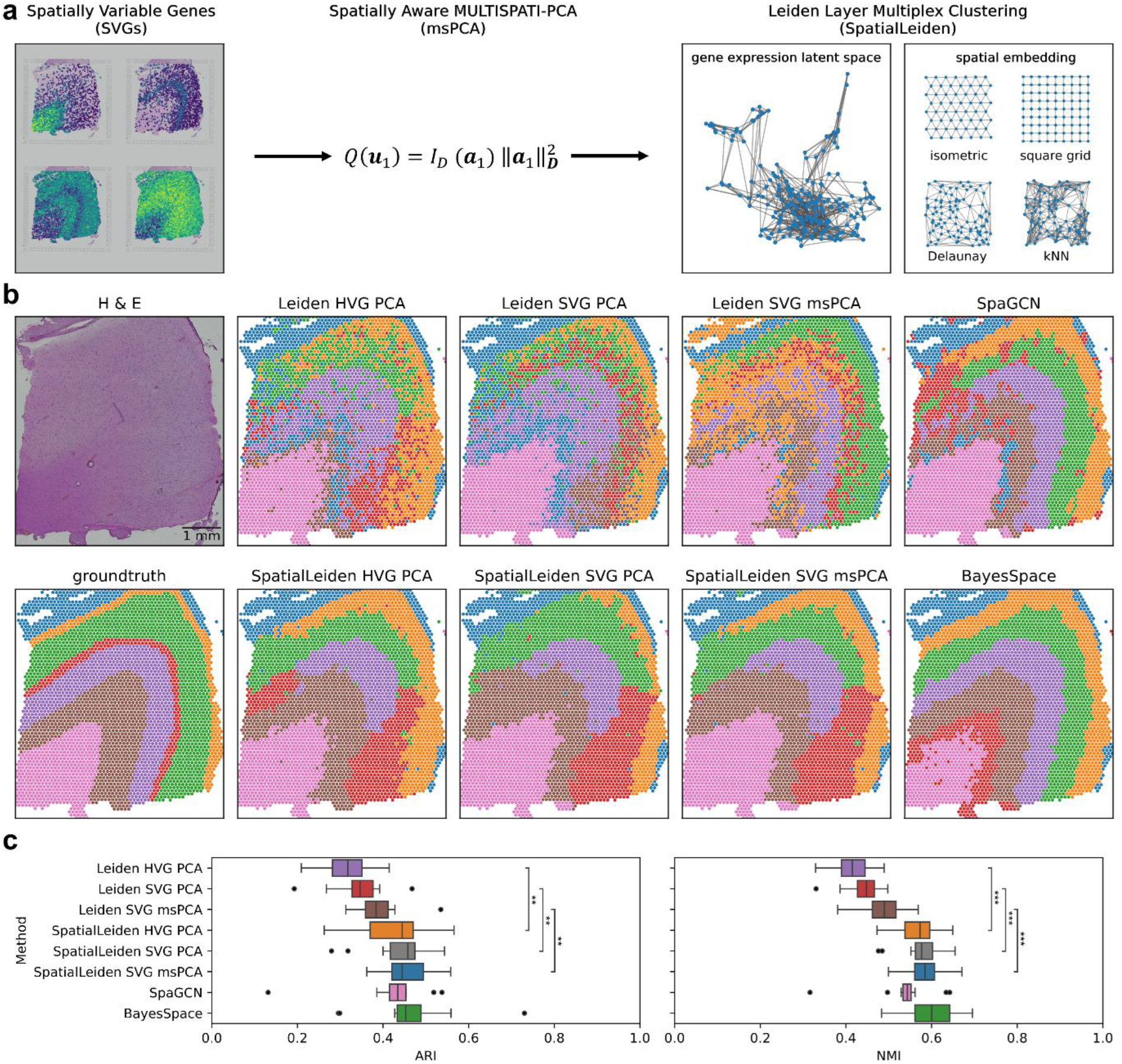
SpatialLeiden workflow. (a) Schematic of data processing and modelling steps to enable spatially-aware Leiden clustering. Feature selection is performed by spatially variable genes (SVGs), dimensionality reduction is performed by a spatially aware MULTISPATI-PCA (msPCA), and clustering is performed by the Leiden layer multiplex algorithm with both gene expression and spatial embeddings (SpatialLeiden). (b) Histology and manually annotated neocortex layered domains for the mouse brain DLPFC (slice 151673) and spatial domains detected by Leiden, SpatialLeiden, SpaGCN and BayesSpace. (c) Boxplot of Adjusted Rand Index (ARI) and Normalised Mutual Information (NMI) for all 12 DLPFC samples. Center line: median; box limits: upper and lower quartiles; whiskers: 1.5× interquartile range; dots: outliers; asterisks: significance (FWER, one-sided Wilcoxon signed-rank test).

While use of SVGs over HVGs yielded only minor improvements, we observed substantial improvement in performance when using spatially aware dimensionality reduction (msPCA) and using SpatialLeiden over non-spatial Leiden, revealing a better representation of the neocortex layering pattern (**Figure 1b**). We quantitatively evaluated performance of the different clustering strategies using the Adjusted Rand Index (ARI) and Normalised Mutual Information (NMI) score, showing significant improvements of SpatialLeiden over the non-spatial Leiden implementation, with performance that was better than SpaGCN and comparable to BayesSpace (**Figure 1 c, Supplemental Figure 1-3, Supplemental table 1-2**) at a fraction of the processing time. SpatialLeiden performed favourably when we compared its performance to other tools in a recent benchmark study, ranking 5^th^ of 15 tools(4), (see **Supplemental Methods** for further details).

As with other multi-modal clustering approaches, careful consideration has to be paid to a number of parameters including the definition and weighting of neighbours across modalities, the resolution to be applied to each modality, and the weight of each modality (**Supplemental Figure 4**). Furthermore, the spatial relationship between cells can be modelled in different ways; a regular grid pattern is suitable for Visium (isometric) and binned Stereo-seq (square), while for imaging-based spatially resolved transcriptomics methods Delaunay triangulation or *k*-nearest neighbours can be used to define the topological layer.

To investigate the performance of SpatialLeiden across technologies, tissues and topological modelling we analysed a number of datasets (Stereo-Seq mouse embryo(10), BaristaSeq mouse brain primary cortex, MERFISH mouse brain hypothalamus preoptic area(11), osmFISH mouse brain somatosensory cortex, STARmap mouse brain medial prefrontal cortex(12,13), STARmap1k mouse brain visual cortex) and demonstrated exceptional improvements over non-spatially aware Leiden clustering. SpatialLeiden ranked as the best performing spatial domain clustering methods for Stereo-Seq, MERFISH and osmFISH, while demonstrating top tier performance for the other datasets (**Figure 2**). For imaging-based spatial transcriptomics methods we found that modelling the topological neighbourhood using the 10 *k*-nearest neighbours generally performed better than using Delaunay triangulation. All non-Stereo-seq dataset were processed within 2 minutes, utilising less than 400 MB of RAM. All Stereo-seq samples were processed within 8 minutes utilising a maximum of 3.5 GB of RAM (**Supplemental Figure 5**).

**Figure 2:**
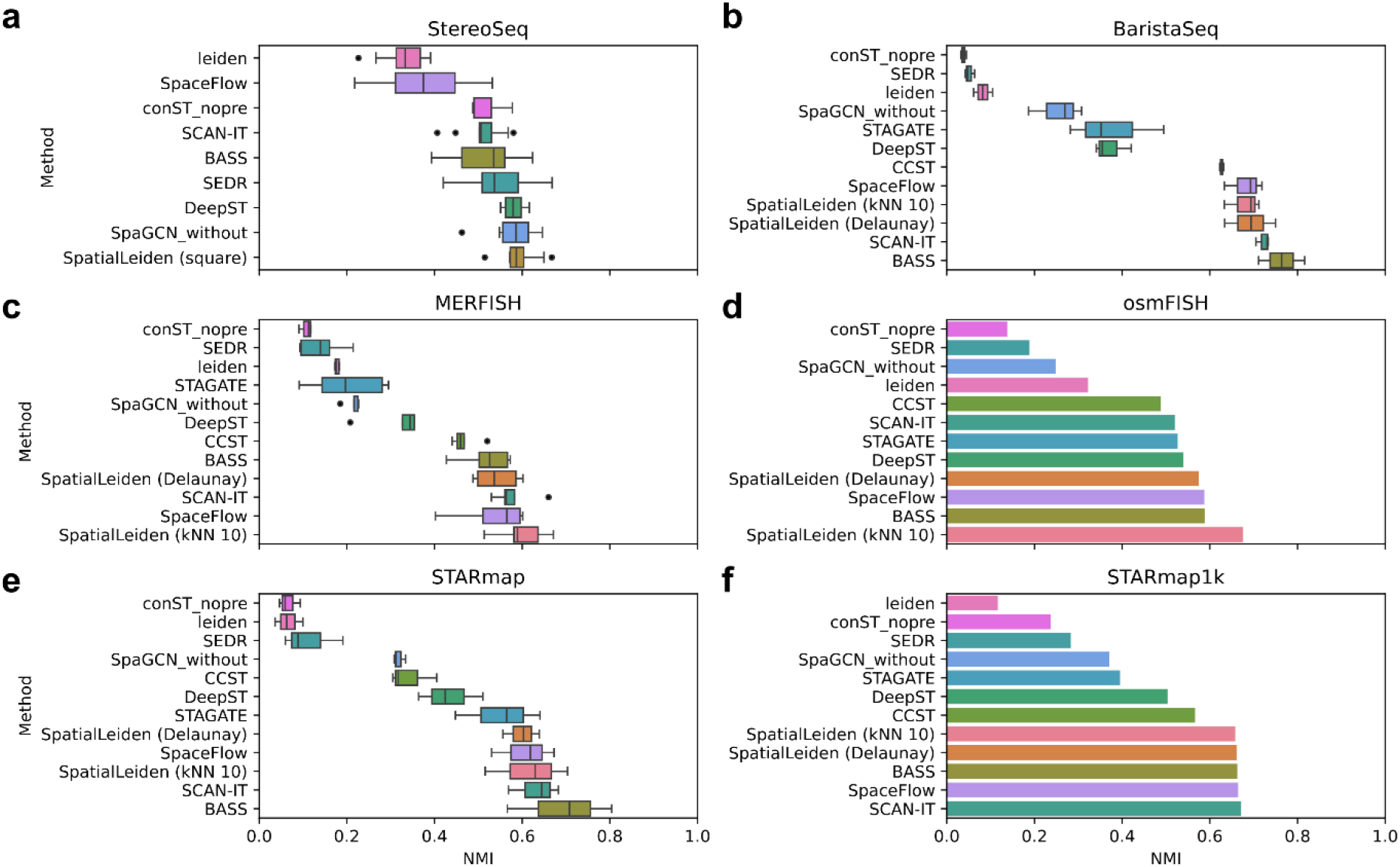
Performance of SpatialLeiden across technologies and tissues: (a) Stereo-Seq of the mouse embryo at various development stages; (b) BaristaSeq of mouse primary cortex; (c) MERFISH mouse brain hypothalamus preoptic area; (d) osmFISH of mouse somatosensory cortex; (e) STARmap mouse brain medial prefrontal cortex; and (f) STARmap* of mouse visual cortex. Performance metrics of other tools are taken from Yuan et al 2024(4). SpatialLeiden was run with 5 different random seeds and median results were reported per sample.

## Conclusions

Our results show that the reference implementation of the Leiden algorithm can indeed be used as a spatially-aware clustering algorithm. Subsequent studies that compare spatially-aware clustering algorithms should clearly state that they compare to non-spatial implementation of Leiden, rather than misclassifying Leiden as a non-spatial algorithm. We describe the different steps at which spatial awareness can be introduced into the analysis, and our implementation allows easy parameterisation of key considerations for modelling gene and spatial modalities. While many spatial domain clustering tools rely on spatially-aware dimensionality reduction approaches this is often followed by non-spatial clustering and we expect these methods to improve with spatially-aware clustering such as SpatialLeiden.

The same way Leiden became the method of choice for clustering of single-cell data, we believe that SpatialLeiden will become the method of choice for spatial data owing to its efficiency, simplicity, and ease of integration into existing analysis workflows.

## Methods

### Data processing

Data was analysed using python (v3.10.14), Scanpy(14) (v1.10.1), and Squidpy(15) (v1.4.1).

### Topological neighbourhood graph generation

The neighbors of each cell were defined depending on the technology. For datasets with a regular grid the neighbors were defined using squidpy.gr.spatial_neighbors with coord_type *‘grid’* and n_neighs set to 6 or 4 for Visium and Stereo-seq, respectively. For all other datasets the neighbors were defined either using Delaunay triangulation (squidpy.gr.spatial_neighbors with delaunay=True) or using the 10 nearest-neighbors (squidpy.gr.spatial_neighbors with coord_type *’generic’* and n_neighs set to 10). The untransformed neighbourhood graph *‘spatial_connectivities’* was used as is for regular grids as all connections are equidistant. For Delaunay triangulation and kNN the *‘spatial_distances’* were transformed to *‘spatial_connectivities’* via the following formula:

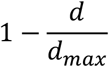

### HVG and SVG detection

Highly variable genes (HVGs) were detected based on filtered count data (scanpy.pp.highly_variable_genes with flavor *‘seurat_v3’*). To detect spatially variable genes (SVGs), first the neighbours were defined as described above and the Moran’s *I* score was calculated for all genes using squidpy.gr.spatial_autocorr with mode *‘moran’* and selecting the top 3,000 scoring genes. Gene selection was only performed for capture-based spatial transcriptomics technologies (Visium, Stereo-seq).

### MULTISPATI-PCA

We implemented MULTISPATI-PCA(6) in python (https://github.com/HiDiHlabs/multiSPAETI v0.1.0) and used it to perform spatially-aware dimensionality reduction. The topological neighbourhood graph *‘spatial_connectivities’* was used to calculate 30 components (corresponding to the 30 largest eigenvalues) based on the 3,000 HVGs / SVGs (Visium, Stereo-seq) or all genes in the case of imaging-based technologies (STARmap, STARmap*, MERFISH, BaristaSeq, osmFISH) as for normal PCA.

### Latent neighbourhood graph generation

The topological neighbourhood graph in the Visium grid *‘spatial_distances’* from squidpy.gr.spatial_neighbors (as described in SVG detection) is used as the spatial layer for Leiden clustering. To build the neighbourhood graph in latent space of gene expression we first calculated the first 30 principal components based on the top 3,000 variable genes (scanpy.tl.pca). We identified the 15 nearest-neighbours per spot (scanpy.pp.neighbors) based on the PCA or MULTISPATI-PCA results from either the HVGs / SVGs (Visium, Stereo-seq) or for all genes in the case of imaging-based technologies (STARmap, STARmap*, MERFISH, BaristaSeq, osmFISH) and used the resulting *‘connectivities’*.

### Non-spatially aware Leiden

We used Leiden(1) (https://github.com/vtraag/leidenalg, v0.10.2) as implemented in Scanpy with the default parameters and varied the resolution to achieve the correct number of clusters for each of the DLPFC datasets following the approach of the SpaGCN.search_res function (https://github.com/jianhuupenn/SpaGCN).

### Spatially-aware Leiden multiplex (SpatialLeiden)

We implemented a spatially aware version of Leiden(https://github.com/HiDiHlabs/SpatialLeiden) by using the Layer multiplex(16). An additional graph encoding for the topological neighbourhood of the cells was added as secondary layer in addition to the layer encoding gene expression in latent space. The additional spatial layer was encoded as RBConfigurationVertexPartition as is the default for the scanpy implementation for the latent space graph. The optimal clustering was identified by running the Optimiser.optimise_partition_multiplex from leidenalg until convergence. As only the ratio of the layer_weights is relevant the weight for the gene expression latent space layer was kept at 1 and the weight for the topological neighbourhood was set depending on technology and method of neighbourhood definition. The resolution for the latent space partition was set by running the standard Leiden clustering and identifying the resolution which yields the correct number of clusters. The resolution of the topological partition was then varied to identify the correct number of clusters in the multiplex Leiden using the same approach as described for the standard Leiden method.

### Implementation and comparison to other spatial clustering algorithms

Implementation and comparison to other spatial clustering algorithms is described in the online methods.

## Abbreviations

(SVGs): Spatially variable genes
(HVGs): Highly variable genes
(msPCA): MUTLISPATI-PCA
(DLPFC): Dorsolateral prefrontal cortex
(ARI): Adjusted Rand Index
(NMI): Normalised Mutual Information

## Declarations

### Ethics approval and consent to participate

Not applicable.

### Consent for publication

Not applicable.

### Availability of data and materials

This study used publicly available spatially resolved transcriptomics data of the mouse brain DLPFC profiled on the 10x Visium platform (http://research.libd.org/spatialLIBD/). Public Stereo-seq, STARmap, STARmap*, MERFISH, osmFISH, and BaristaSeq datasets were downloaded from http://sdmbench.drai.cn/. Performance metrics of other tools presented in Figure 2 were taken from Yuan et al(4). We also provide a repository with Jupyter Notebooks for reproducing all results and figures of this study https://github.com/HiDiHlabs/SpatialLeiden-Study.

### Authors’ contributions

NI conceived and designed the study. SS implemented the spatially-aware Leiden. NMB, SS implemented the SpatialLeiden package. NMB implemented MULTISPATI-PCA in python.

SS performed code review. NMB performed data analysis. NI, NMB interpreted and analysed results. NI, NMB, SS, RE proofread and corrected the manuscript. All authors contributed to the article and approved the submitted version.

## Acknowledgements

We thank organisers and participants of the de.NBI BioHackathon SpaceHack 2.0 project in Bielefeld, Germany in December 2023 (**Supplemental Table 3**).

