## Supplement for "SpatialLeiden - Spatially-aware Leiden clustering"

### Supplemental Information

Supplemental information for “SpatialLeiden - Spatially-aware Leiden clustering” by Niklas Müller-Bötticher, Shashwat Sahay, Roland Eils, and Naveed Ishaque.

**Content**

- Supplemental Methods
- Supplemental References
- Supplemental Figures 1-5
- Supplemental Tables 1-3

### Supplemental Methods

#### Data processing

Data was analysed using python (v3.10.14), Scanpy(1) (v1.10.1), and Squidpy(2) (v1.4.1).

#### BayesSpace

BayesSpace(3) is a method that applies Bayesian statistics to a low-dimensional representation of gene expression for spatial clustering, optimising for neighbouring points to having the same cluster membership through a spatial prior. BayesSpace is one of the early and highly cited tools for spatial domain identification. We used BayesSpace (v1.10.1, R v4.3.1) using preprocessing and parameters as recommended in the original publication and package vignette. Our observed results (median ARI: 0.45) were not lower than the recently reported ARIs by Hu et al(4) (0.37).

#### SpaGCN

SpaGCN is a method that utilises a graph convolutional neural network graph to integrates gene expression, spatial location and histology for domain identification. SpaGCN is another one of the early and highly cited tools for spatial domain identification. We used SpaGCN (v1.2.7, python v3.8.18) using preprocessing and parameters as defined in the original publication and package tutorial. Our observed results (median ARI: 0.43) were in line with the recently reported ARIs by Hu et al(4) (0.41).

#### ARI and NMI calculation

The Adjusted Rand Index (ARI) and Normalized Mutual Information (NMI) were calculated using the adjusted_rand_score and normalized_mutual_info_score functions in scikit-learn(5) (v1.5.0) with all parameters at default values.

#### Data analysis

##### Capture-based technologies

Capture-based datasets (Visium human brain dorsolateral prefrontal cortex, Stereo-Seq mouse embryo) were preprocessed by removing genes that appear in less than 3 spots (scanpy.pp.filter_genes), normalizing the total count of each cell (scanpy.pp.normalize_total with target_sum 1e4), and log transforming the data (scanpy.pp.log1p). The layer weight ratio of topological to latent space graph was set to 0.8 in SpatialLeiden unless indicated otherwise.

SpatialLeiden demonstrated significantly improved ARI and NMI over the non-spatial Leiden clustering for the Visium DLPFC dataset (ARI-HVG p=0.0007 FWER=0.0022, ARI-SVG p=0.0017 FWER=0.0051, ARI-SVG-msPCA=0.0007 FWER=0.0022, NMI-HVG p=0.00024 FWER=0.00073, NMI-SVG p=0.00024 FWER=0.00073, NMI-SVG-msPCA=0.00024 FWER=0.00073, one-sided Wilcoxon signed-rank test and Bonferroni corrected family-wise error rate, **Figure 1c**).

##### Imaging-based technologies

Imaging-based datasets (MERFISH mouse brain hypothalamus preoptic area, STARmap mouse brain medial prefrontal cortex, STARmap* mouse visual cortex, BaristaSeq mouse primary cortex, osmFISH mouse somatosensory cortex) were log transformed (scanpy.pp.log1p). The layer weight ratio for SpatialLeiden for MERFISH was set to 1 and 1.8 for Delaunay triangulation and kNN (10 neighbors), respectively. STARmap: 1 and 1.6, STARmap*: 0.8 and 1.4, BaristaSeq: 1.2 and 1.8, osmFISH: 0.8 and 1.2.

#### Runtime performance metrics

To calculate runtime performance metrics jobs were submitted using slurm sbatch to a single compute node (-N 1) with 8 CPUs (-n 8) with each node consisting of a Dell PowerEdge C6520 with 2x Intel Xeon Gold 6130 or 6252 @ 2.1 GHz. The metrics were calculated from the slurm job information (using *'Elapsed'* as wall time, *'TotalCPU'* as CPU time, and *'MaxRSS'* as maximum memory usage) for the entire workflow including data loading, preprocessing, dimensionality reduction, and clustering.

### Comparison to other benchmarking studies

We compared the obtained median NMI (0.58) for multiplex Leiden clustering using SVGs and MULTISPATI-PCA on the Visium DLPFC dataset (**Figure 1, Supplementary Tables 1,**

**2**) to those in a study by Yuan et al(6). They reported the NMI across 10 replicate runs for each sample over 14 tools in their Source Data Figure 2. The median NMI per tool, ordered by NMI, were: Louvain 0.24; Leiden 0.25; SpaGCN (w/ H&E): 0.48; SpaceFlow 0.49; STAGATE 0.50; CCST 0.51; conST_nopre 0.52; SpaGCN (w/o H&E): 0.53; SEDR 0.53; stLearn 0.54; BayesSpace 0.60; SCAN-IT 0.61; BASS 0.61; DeepST 0.62.

### Supplemental Figure 1

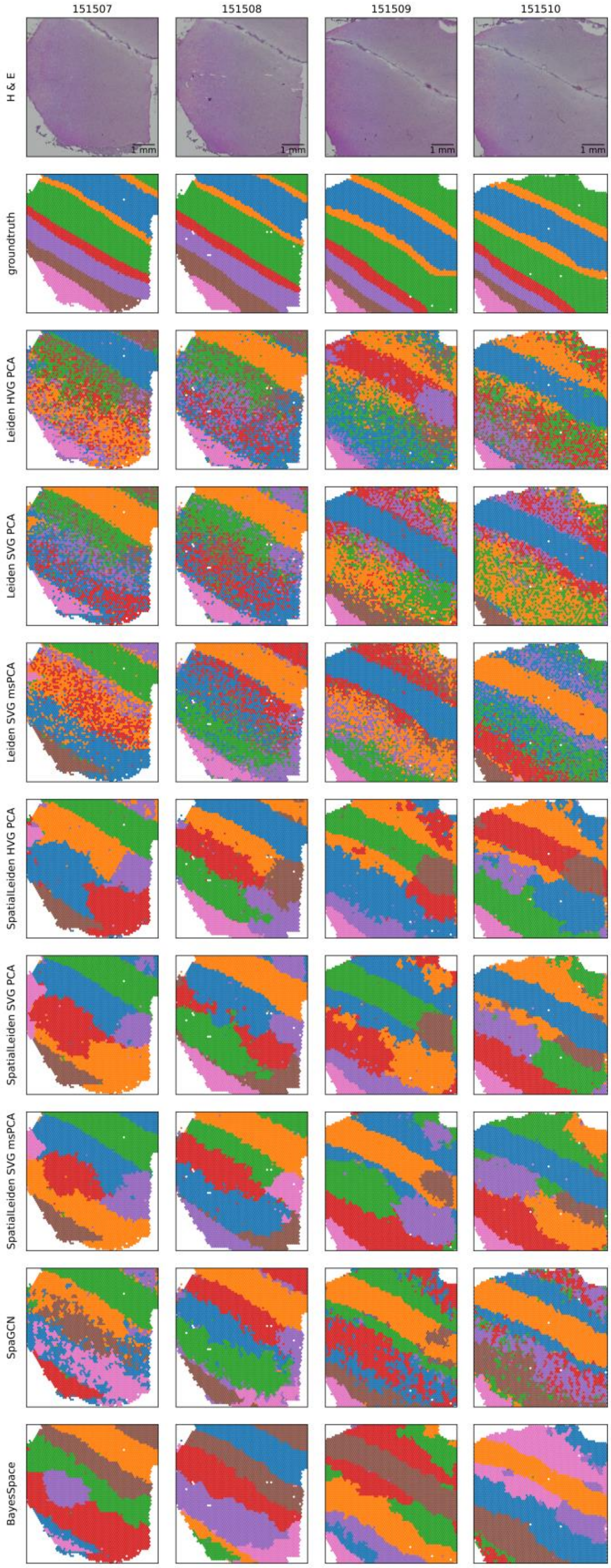


**Supplemental Figure 1**: Clustering results obtained from samples from Br5292.

### Supplemental Figure 2

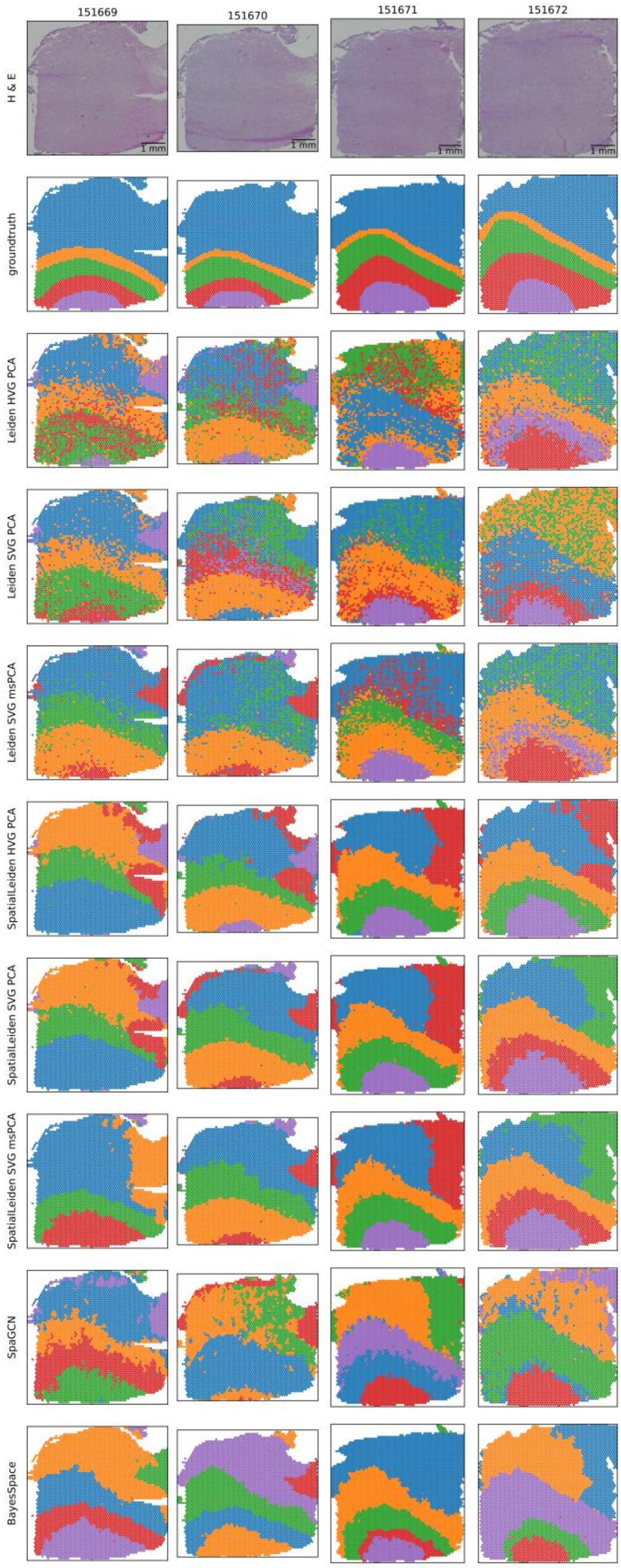


**Supplemental Figure 2**: Clustering results obtained from samples from Br5295.

### Supplemental Figure 3

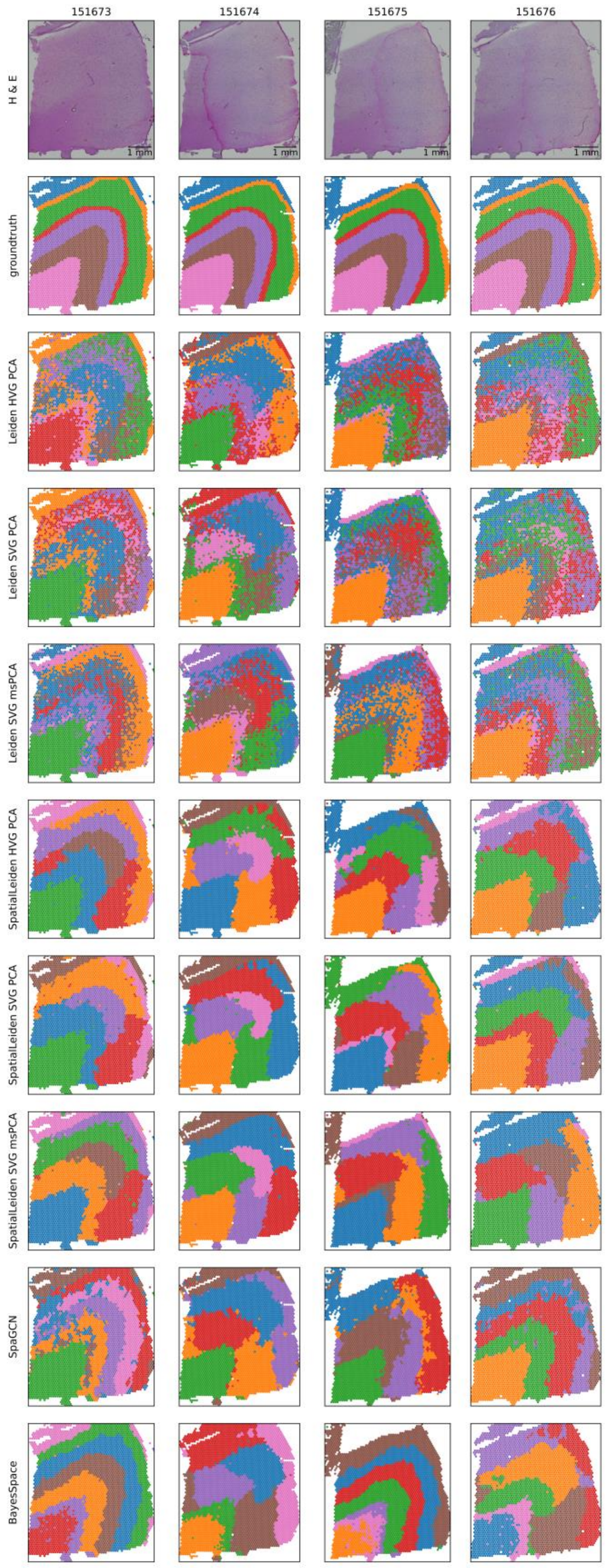


**Supplemental Figure 3**: Clustering results obtained from samples from Br8100.

### Supplemental Figure 4

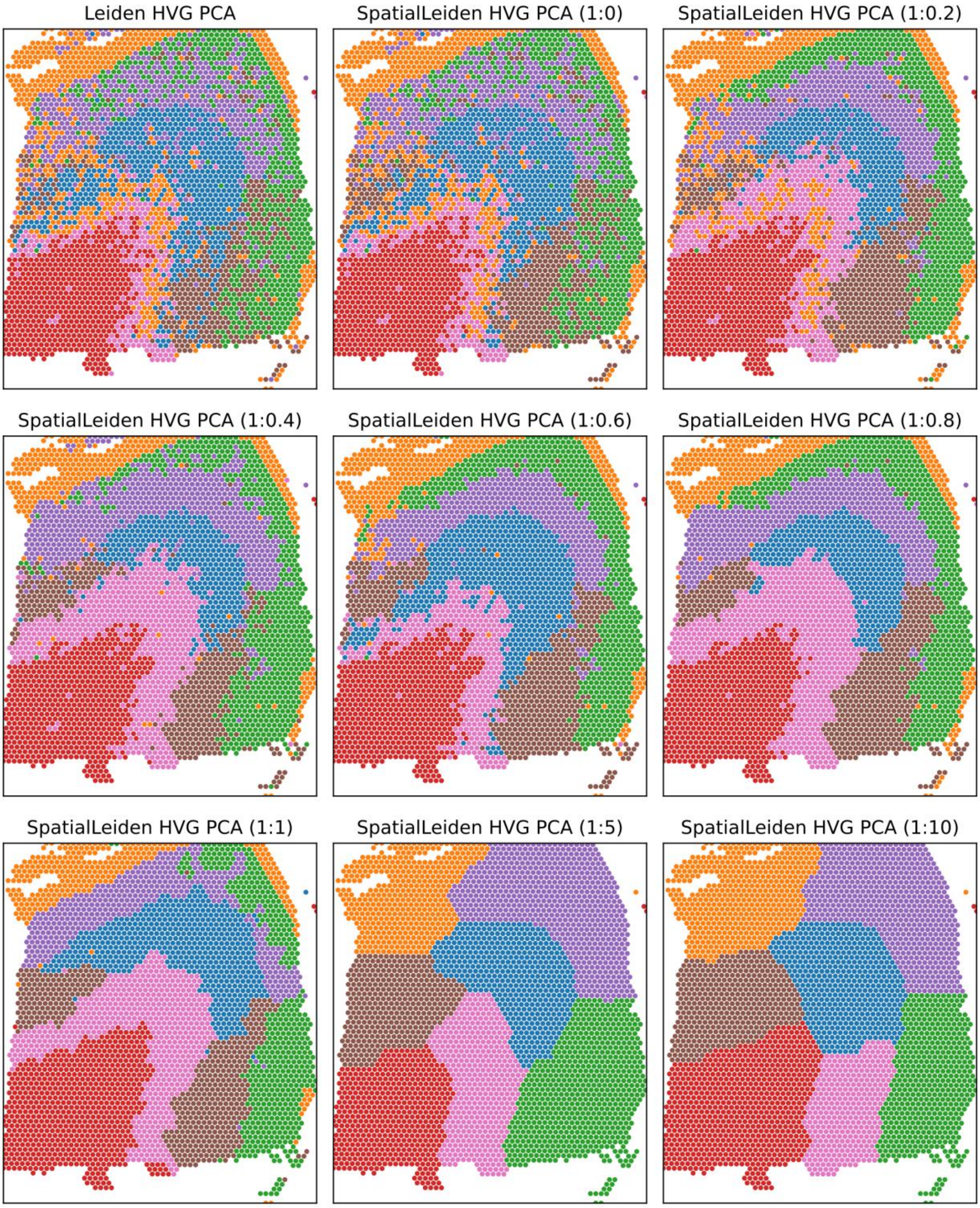


**Supplemental Figure 4**: The effect of weighting the gene expression vs spatial layers. Shown are the clustering results for sample Br8100-151673 using non-spatial Leiden and SpatialLeiden. The ratio of weights (expression:spatial) is shown in parenthesis in the subplot titles.

### Supplemental Figure 5

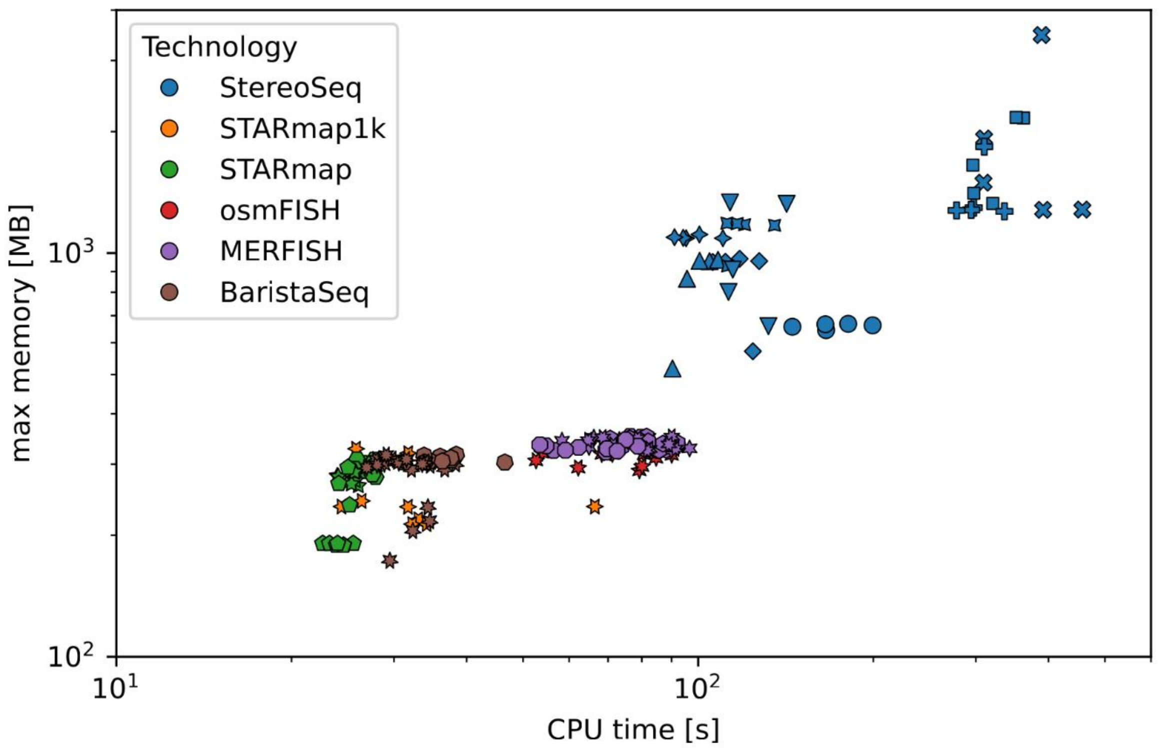


**Supplemental Figure 5**: Runtime statistics. Scatter plot of total CPU time vs max memory used to process datasets across technologies. Colours represent the technology and the shape encodes the specific sample for each dataset. Each sample was processed 5 times with different starting seeds (0, 1, 2, 3, 4).

### Supplemental Table 1: ARI for Visium DLPFC

| Sample | BayesSpace | Leiden HVG PCA | Leiden SVG PCA | Leiden SVG msPCA | SpaGCN | SpatialLeiden HVG PCA | SpatialLeiden SVG PCA | SpatialLeiden SVG msPCA |
|---|---|---|---|---|---|---|---|---|
| **Br5292_151507** | 0.477 | 0.329 | 0.347 | 0.409 | 0.423 | 0.439 | 0.460 | **0.506** |
| **Br5292_151508** | **0.479** | 0.306 | 0.336 | 0.379 | 0.436 | 0.450 | 0.456 | 0.440 |
| **Br5292_151509** | **0.447** | 0.259 | 0.392 | 0.428 | 0.422 | 0.371 | 0.400 | 0.399 |
| **Br5292_151510** | 0.442 | 0.360 | 0.346 | 0.416 | 0.453 | 0.379 | **0.469** | 0.449 |
| **Br5595_151669** | 0.459 | 0.290 | 0.374 | 0.376 | 0.394 | 0.262 | 0.280 | **0.477** |
| **Br5595_151670** | **0.427** | 0.209 | 0.193 | 0.313 | 0.131 | 0.315 | 0.318 | 0.375 |
| **Br5595_151671** | **0.730** | 0.252 | 0.468 | 0.535 | 0.538 | 0.523 | 0.542 | 0.558 |
| **Br5595_151672** | 0.435 | 0.414 | 0.384 | 0.410 | 0.453 | **0.566** | 0.544 | 0.546 |
| **Br8100_151673** | **0.559** | 0.338 | 0.303 | 0.352 | 0.439 | 0.496 | 0.436 | 0.428 |
| **Br8100_151674** | 0.296 | 0.369 | 0.365 | 0.388 | 0.385 | 0.461 | **0.491** | 0.490 |
| **Br8100_151675** | **0.516** | 0.348 | 0.343 | 0.361 | 0.432 | 0.456 | 0.422 | 0.439 |
| **Br8100_151676** | 0.299 | 0.292 | 0.267 | 0.323 | **0.518** | 0.367 | 0.466 | 0.362 |
| **Median** | 0.453 | 0.317 | 0.346 | 0.384 | 0.434 | 0.445 | **0.458** | 0.445 |

### Supplemental Table 2: NMI for Visium DLPFC

| Sample | BayesSpace | SpaGCN | Leiden HVG PCA | Leiden SVG PCA | Leiden SVG msPCA | SpatialLeiden HVG PCA | SpatialLeiden SVG PCA | SpatialLeiden SVG msPCA |
|---|---|---|---|---|---|---|---|---|
| **Br5292_151507** | **0.630** | 0.529 | 0.408 | 0.452 | 0.525 | 0.547 | 0.572 | 0.618 |
| **Br5292_151508** | **0.616** | 0.545 | 0.402 | 0.422 | 0.462 | 0.585 | 0.568 | 0.573 |
| **Br5292_151509** | **0.609** | 0.542 | 0.429 | 0.491 | 0.514 | 0.578 | 0.586 | 0.583 |
| **Br5292_151510** | 0.567 | 0.546 | 0.422 | 0.439 | 0.501 | 0.563 | **0.607** | 0.568 |
| **Br5595_151669** | 0.591 | 0.535 | 0.346 | 0.458 | 0.462 | 0.473 | 0.485 | **0.601** |
| **Br5595_151670** | **0.545** | 0.316 | 0.329 | 0.330 | 0.381 | 0.476 | 0.476 | 0.499 |
| **Br5595_151671** | **0.682** | 0.642 | 0.386 | 0.498 | 0.569 | 0.639 | 0.654 | 0.671 |
| **Br5595_151672** | 0.590 | 0.549 | 0.489 | 0.491 | 0.525 | 0.649 | 0.655 | **0.656** |
| **Br8100_151673** | **0.695** | 0.561 | 0.459 | 0.443 | 0.496 | 0.628 | 0.566 | 0.586 |
| **Br8100_151674** | 0.483 | 0.497 | 0.459 | 0.456 | 0.484 | 0.568 | 0.601 | **0.603** |
| **Br8100_151675** | **0.681** | 0.541 | 0.440 | 0.429 | 0.461 | 0.583 | 0.552 | 0.540 |
| **Br8100_151676** | 0.496 | **0.634** | 0.391 | 0.386 | 0.440 | 0.509 | 0.581 | 0.501 |
| **Median** | **0.600** | 0.543 | 0.415 | 0.448 | 0.490 | 0.573 | 0.577 | 0.585 |

### Supplemental Table 3: SpaceHack v2.0 participants

| Name | Email | ORCiD | Affiliation |
|---|---|---|---|
| Ahmed Mahfouz | | 0000-0001-8601-2149 | Department of Human Genetics, Leiden University Medical Center, Einthovenweg 20, 2333 ZC Leiden, Netherlands |
| Alexander Kanitz | | 0000-0002-3468-0652 | Biozentrum, University of Basel, Spitalstrasse 41, 4056 Basel, Switzerland |
| Anastasiia Okhtienko | | 0009-0003-5886-811X | Technical University of Munich, School of Medicine, Institute of Virology, Munich, Germany |
| Brian Long | | 0000-0002-7793-5969 | Allen Institute for Brain Science, 615 Westlake Ave N, Seattle, WA, USA 98109 |
| Charlotte Soneson | | 0000-0003-3833-2169 | Friedrich Miescher Institute for Biomedical Research, Basel, Switzerland |
| Christoph Kuppe | | 0000-0003-4597-9833 | Division of Nephrology and Clinical Immunology, RWTH Aachen University, Aachen, Germany |
| Daryna Pikulska | | 0009-0005-1638-0268 | Department of Nephrology, Rheumatology, and Clinical |

| | | | |
|---|---|---|---|
| | | | Immunology, University Hospital RWTH Aachen, Pauwelsstraße 30, Aachen, Germany |
| Divya Sitani | | 0000-0001-5138-6108 | Data Science Lab, Hertie School, 10117 Berlin, Germany |
| Estella Yixing Dong | | 0009-0003-5115-5686 | Biomedical Data Science Center, Centre hospitalier universitaire vaudois, Rue du Bugnon 21, 1011 Lausanne |
| Fadhl Alakwaa | | 0000-0001-5349-7960 | Department of Internal Medicine, Division of Nephrology, University of Michigan, Ann Arbor, Michigan, USA |
| Florian Heyl | | 0000-0002-3651-5685 | German Cancer Research Center (DKFZ), Division of Computational Genomics and Systems Genetics, Im Neuenheimer Feld 280, 69120 Heidelberg, Germany |
| Francesca Antonella Luongo | | 0009-0005-6475-9029 | ETHZ, Rämistrasse 101, 8092 Zürich / CSEM Hegenheimermattweg 167A, 4123 Allschwil |
| Georgios Gavriilidis | | 0000-0003-2575-4354 | Institute of Applied Biosciences, Centre for |

| | | | |
|---|---|---|---|
| | | | Research and Technology Hellas, Thessaloniki, Greece |
| Giorgia Moranzoni | | 0000-0001-8065-6277 | DTU Health Tech, Technical University of Denmark, Ørsteds Plads, Building 345C, 2800, Kgs. Lyngby, Denmark |
| Jieran Sun | | 0000-0002-7996-3840 | Biomedical Data Science Center, Centre hospitalier universitaire vaudois, Rue du Bugnon 21, 1011 Lausanne |
| Kim Vucinic | | 0009-0008-1553-1935 | Laboratory of Genomics and Bioinformatics, Institute of Molecular Genetics of the Czech Academy of Sciences, 142 20, Prague 4, Czech Republic |
| Kirti Biharie | | 0000-0002-8274-8439 | Delft Bioinformatics Lab, Delft University of Technology, Van Mourik Broekmanweg 6, 2628 XE Delft, Netherlands |
| Lena Perry | | | University of Michigan, UROP(Undergraduate Research Opportunity Program), Ann Arbor, Michigan |
| Liya Zaygerma n | | 0009-0005-4947-6223 | Helmholtz Zentrum München, Institute of Computational Biology, Ingolstädter |

| | | | |
|---|---|---|---|
| | | | Landstraße 1 · D-85764 Neuherberg |
| Louis Kümmerle | | 0000-0002-9193-1243 | Helmholtz Zentrum München, Institute of Computational Biology, Ingolstädter Landstraße 1 · D-85764 Neuherberg |
| Lucie Pfeiferova | | 0000-0003-1089-0329 | Laboratory of Genomics and Bioinformatics, Institute of Molecular Genetics of the Czech Academy of Sciences, 142 20, Prague 4, Czech Republic. |
| Mar Muniz Moreno | | 0000-0002-2662-890X | OMAPiX GmbH Raiffeisenstraße 3 70764 Langenfeld Germany |
| Marco Varrone | | 0000-0002-0538-3464 | Department of Computational Biology, University of Lausanne, Lausanne, Switzerland |
| María Calleja C | | 0000-0003-0688-0793 | CIMA Universidad de Navarra, Pamplona, Spain |
| Mark D. Robinson | | 0000-0002-3048-5518 | SIB Swiss Institute of Bioinformatics and Department of Molecular Life Sciences, University of Zurich, |

| | | | |
|---|---|---|---|
| | | | Winterthurerstrasse 190, 8057 Zurich, Switzerland |
| Meghan A. Turner | | 0000-0003-2451-5036 | Allen Institute for Brain Science, 615 Westlake Ave N, Seattle, WA 98109 |
| Michael Fletcher | | 0000-0003-1589-7087 | Nature Genetics, Heidelberger Platz 3, 10715 Berlin, Germany |
| Naveed Ishaque | | 0000-0002-8426-901X | Berlin Institute of Health at Charité Berlin, Anna-Louisa-Karsch-Straße 2, 10178 Berlin |
| Nigel Shijie Chou | | 0000-0002-6280-7729 | Genome Institute of Singapore, Agency for Science, Technology and Research (A*STAR), Singapore, Singapore |
| Niklas Müller-Bötticher | | 0000-0001-5103-7282 | Berlin Institute of Health at Charité Berlin, Anna-Louisa-Karsch-Straße 2, 10178 Berlin |
| Paul Kiessling | | 0000-0002-9794-9532 | Department of Nephrology, Rheumatology, and Clinical Immunology, University Hospital RWTH Aachen, Pauwelsstraße 30, Aachen, Germany |
| Peiying Cai | | 0009-0001-9229-2244 | Department of Molecular Life Sciences, University of Zurich, |

| | | | |
|---|---|---|---|
| | | | Winterthurerstrasse 190, 8057 Zurich, Switzerland |
| Qirong Mao | | 0000-0001-7938-6660 | Department of Human Genetics, Leiden University Medical Center, Einthovenweg 20, 2333 ZC Leiden, Netherlands |
| Rasool Saghaleyni | | 0000-0003-0956-039X | Department of Biology and Biological Engineering, Chalmers University of Technology, 412 96 Gothenburg, Sweden |
| Sebastian Tiesmeyer | | 0000-0002-0492-1523 | Berlin Institute of Health at Charité Berlin, Anna-Louisa-Karsch-Straße 2, 10178 Berlin |
| Shahul Alam | | 0000-0002-4821-7079 | Computational Biology Department, Carnegie Mellon University, 5000 Forbes Avenue, Pittsburgh, PA 15232 |
| Siao-Han Wong | | 0000-0001-6429-2880 | German Cancer Research Center (DKFZ), Im Neuenheimer Feld 280, 69120 Heidelberg, Germany |
| Sikander Hayat | | 0000-0001-5919-8371 | University Hospital RWTH Aachen, Pauwelsstraße 30, Aachen, Germany |

| | | | |
|---|---|---|---|
| Søren Helweg Dam | | 0000-0003-0755-0016 | DTU Health Tech, Technical University of Denmark, Ørsteds Plads, Building 345C, 2800, Kgs. Lyngby, Denmark |
| Sven Twardziok | | 0000-0002-0326-5704 | Berlin Institute of Health at Charité Berlin, Anna-Louisa-Karsch-Straße 2, 10178 Berlin |
| Teresa Zulueta-Coarasa | | 0000-0002-0456-6912 | European Molecular Biology Laboratory, European Bioinformatics Institute, Hinxton, United Kingdom |
| Thomas Chartrand | | 0000-0002-7093-8681 | Allen Institute for Neural Dynamics, 615 Westlake Ave N, Seattle, WA 98109 |
| Yuzhou Chang | | 0000-0003-4893-1886 | Department of Biomedical Informatics, College of Medicine, Ohio State University, Columbus, OH 43210, USA. |
| Zaira Seferbekova | | 0000-0002-3131-3697 | Artificial Intelligence in Oncology (B450), Deutsches Krebsforschungszentrum, Im Neuenheimer Feld 280, 69120 Heidelberg |
